# Electrophysiological and Transcriptomic Features Reveal a Circular Taxonomy of Cortical Neurons

**DOI:** 10.1101/2021.03.24.436849

**Authors:** Alejandro Rodríguez-Collado, Cristina Rueda

## Abstract

The complete understanding of the mammalian brain requires exact knowledge of the function of each of the neurons composing its parts. To achieve this goal, an exhaustive, precise, reproducible, and robust neuronal taxonomy should be defined. In this paper, a new circular taxonomy based on transcriptomic features and novel electrophysiological features is proposed. The approach is validated by analysing more than 1850 electrophysiological signals of different mouse visual cortex neurons proceeding from the Allen Cell Types Database.

The study is conducted on two different levels: neurons and their cell-type aggregation into Cre Lines. At the neuronal level, electrophysiological features have been extracted with a promising model that has already proved its worth in neuronal dynamics. At the Cre Line level, electrophysiological and transcriptomic features are joined on cell types with available genetic information. A taxonomy with a circular order is revealed by a simple transformation of the first two principal components that allow the characterization of the different Cre Lines. Moreover, the proposed methodology locates other Cre Lines in the taxonomy that do not have transcriptomic features available. Finally, the taxonomy is validated by Machine Learning methods which are able to discriminate the different neuron types with the proposed electrophysiological features.

## 1 INTRODUCTION

Understanding the nervous system’s mechanisms and capabilities, such as the conscience and cognition, remain one of the most challenging and interesting unresolved problems in biology. It requires a precise description of the structure and function of each brain region, including the study of the neuronal circuits and neurons composing them. Furthermore, one fundamental prerequisite in this subsequent structural division study is the creation of a solid neuronal cell-type classification or taxonomy. As (Zeng and Sanes, 2017) explain, cells in the nervous system should be hierarchically classified into different levels, mainly into classes, subclasses and types. This property makes the taxonomy define relationships between cell types, as well as making it easier to update in the light of new information. At class level, cells are classified into non-neuronal cells and neurons, which in turn can be classified into excitatory or pyramidal neurons and inhibitory neurons or interneurons. Excitatory neurons are habitually morphologically spiny, have a long apical dendrite and exhibit less variability in their electrophysiological features. Inhibitory neurons are broadly aspiny, have a more compact dendritic structure and tend to spike faster. Also, at class level, neurons can be classified based on their neurotransmitter into GABAergic and glutamatergic. The latter are mostly excitatory and brain-area specific, while the former are broadly inhibitory and not-area specific. According to (Tremblay et al., 2016), at subclass level, GABAergic neurons belong to four broad classes based on the expressed Cre Lines: Pvalb (Parvalbumin) positive, Vip (Vasoactive intestinal peptide) positive, Sst (Somatostin) positive and cells that express 5-hydroxytryptamine receptor 3A (Htr3a) but are Vip negative. On the other hand, glutamatergic neurons belong to different subclasses according to their laminar locations and the locations to which they project their axons. Finally, neurons belong to different types according to the expressed Cre Line. Aside from these previous statements, the different studies unearth discrepancies in terms of number of neuronal types, their characteristics and the existing order between them as is reviewed in the next paragraph.

The definition of a solid neuronal taxonomy is a challenging task. Heterogeneity between cells arises due to different electrophysiological, morphological and/or genetic features, but also due to differences in cell age, environmental conditions and other sources of noise. Another concerning issue is the reproducibility of the approach. The open challenge of creating a neuronal taxonomy has recently generated many studies, mainly due to the increase in data availability as well as the rise in data computational methods. A recent overview of the matter can be found in (Zeng and Sanes, 2017). In particular, the taxonomy of the mouse visual cortex cells has been the focus of recent research. In (Tasic et al., 2016) and (Tasic et al., 2018), taxonomies based on transcriptomic characteristics obtained from single RNA sequencing are presented. Electrophysiological taxonomies are predominantly based on patch-clamp recordings of neuron membrane potential signals that contain Action Potential Curves (APs), as is done in (Ghaderi et al., 2018) and (Teeter et al., 2018). Furthermore (Gouwens et al., 2019) presents a taxonomy based on the combination of electrophysiological and morphological features, while part of this taxonomy is expanded with transcriptomic features in (Gouwens et al., 2020).

It should be noted that electrophysiological features are easier to measure than other types of features and can be simultaneously recorded on hundreds of cells using scalable techniques such as optical imaging of electrical activity (Zeng and Sanes, 2017). Furthermore, taxonomies based on this type of features are easier to reproduce.However, the features traditionally used in this type of taxonomy lack interpretability as they are not directly related to the observed potential difference signal. Most of these studies extract the features with dimensional reducing techniques (as is the case of Ghaderi et al. (2018) or Gouwens et al. (2019) among others) or the features are model parameters such as the Leaky Integrate and Fire (LIF) models (as in Teeter et al. (2018)). A brief overview of the latter models can be found in (Lynch and Houghton, 2015).

The aim of this paper is to derive an electrophysiological-transcriptomic circular taxonomy of visual cortex Cre Lines, using data from *Mus Musculus* of the Allen Cell Types Database (ACTD) (http://celltypes.brain.map.org). This database is freely available and has been the reference data for many authors, such as (Teeter et al., 2018) and references therein. On the one hand, the electrophysiological features at Cre Line level are the median of those generated at the cell level from fitting a Frequency Modulated Möbius (FMM) model to the observed cell signals. The FMM model is a flexible model defined by 13 parameters that accurately describes the AP shape. The monocomponent FMM model is presented in (Rueda et al., 2019) and model extensions to analyse neuronal dynamics are shown in (Rueda et al., 2021) and (Rodríguez-Collado and Rueda, 2021). Some relevant and robust theoretical properties of the model are shown in the former paper while, in the latter, an FMM representation of the famed Hodgkin-Huxley model is proposed.

On the other hand, the observed frequencies of the Cre Types by Cre Line from (Tasic et al., 2016) have been used as transcriptomic features. Finally, the morphological features have not been used as they are sparsely available compared to other kinds of measurements and they seem to be not as discriminant for Cre Lines as the other kinds of features, as seen in (Gouwens et al., 2020).

As experts on *Feature Engineering* state, the expected results in a classification problem are largely dependent on whether useful and discriminant features have been extracted ((Heaton, 2017) and (Duboue, 2020) among recent papers on the matter). In our case, it is shown that the FMM model parameters successfully discriminate signals from different neuronal cell types, which validates the proposed taxonomy.

## 2 MATERIALS AND METHODS

### 2.1 Statistical Methods

Let *X*(*t*_*i*_) denote the potential difference in the neuron’s membrane at each of the observed time points *t*_*i*_, *i* = 1, …, *n*. The latter are assumed to be in [0, 2*π*). Otherwise, consider *t′* ∈ [*t*_0_, *T* + *t*_0_] with *t*_0_ as the initial time value and *T* as the period. Transform the time points by 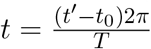.

In this section, the statistical methods used in the manuscript are described. These include the FMM model, Circular Principal Components Analysis (CPCA) and, Machine Learning Supervised Methods.

#### 2.1.1 FMM Model

The proposed model to analyse AP data is a three-component FMM model, as defined in (Rueda et al., 2021) which implies that each AP is modelled using three waves, labeled *A, B* and *C*. The physiological meaning of these waves is given below, after the mathematical presentation.

Mathematically, the waves are defined as follows:

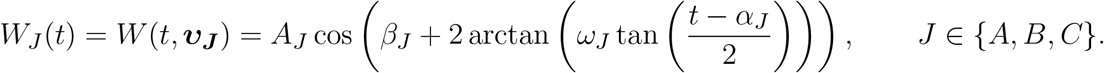

Where ***v***_***J***_ = (*A*_*J*_, *α*_*J*_, *β*_*J*_, *ω*_*J*_) ′ is a four-dimensional parameter describing the shape of the wave. The *A* parameter represents the wave amplitude whereas *α* is a location parameter. The parameters *β* and *ω* determine the skewness and kurtosis of the wave. More details about the interpretation of the parameters can be found in (Rueda et al., 2019).

Moreover, the FMM model is defined as a signal plus error model, as follows:

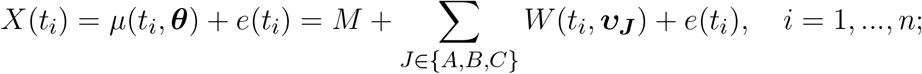

where,

- *θ* = (*M, v*_*A*_, *v*_*B*_, *v*_*C*_) verifying:
  1. *M* ∈ℜ; *v*_*J*_ ∈ ℜ^+^ × [0, 2*π*] × [0, 2*π*] × [0, 1]; *J* ∈ {*A, B, C*}
  2. *α*_*A*_ ≤ *α*_*B*_ ≤ *α*_*C*_
- (*e*(*t*_1_), …, *e*(*t*_*n*_)) ′ ∼ *N*_*n*_(0, *σ*^2^***I***).

The restrictions on the *α*s guarantee identifiability.

Other important parameters of practical use are peak and trough times, denoted by 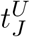 and 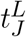 respectively, and the distances between the model waves, denoted by *d*_*JK*_. All of them are defined as follows:

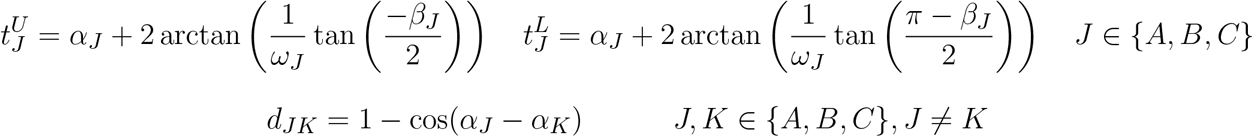

The papers (Rueda et al., 2021) and (Rodríguez-Collado and Rueda, 2021) provide model properties as well as detail the algorithm used to fit the models. In particular, the second paper presents a restricted FMM model for AP trains, while in the first one data from ACTD is concisely analysed. Also, the associated phase space of the model is studied and relevant properties are provided.

In Figure 1, the fitted FMM model prediction and wave decomposition of a GABAergic AP (Figure 1A) and a glutamatergic AP (Figure 1B) are shown. *W*_*A*_ represents the repolarization and, partly, the depolarization while *W*_*B*_ describes the end of the depolarization, and the hyperpolarization. Glutamatergic cells tend to have wider APs with a bigger amplitude (values of *β*_*A*_ smaller than *π* and higher values of *ω*_*A*_ and *A*_*A*_) than GABAergic cells. Furthermore interesting differences can be observed between the two types in terms of the parameters of *W*_*B*_, particularly in *β*_*B*_ and *ω*_*B*_. The third wave, *W*_*C*_, is heteromorphous: in some cases, this wave completes the AP shape (as is typical in GABAergic neurons), while in other cases it accounts for potential differences before and after the spike (as happens in most of the glutamatergic neurons). Also, *W*_*A*_, *W*_*B*_ and *W*_*C*_ seem to be related to the potassium, sodium and calcium conductances that appear in (Gouwens et al., 2018).

**Figure 1.**
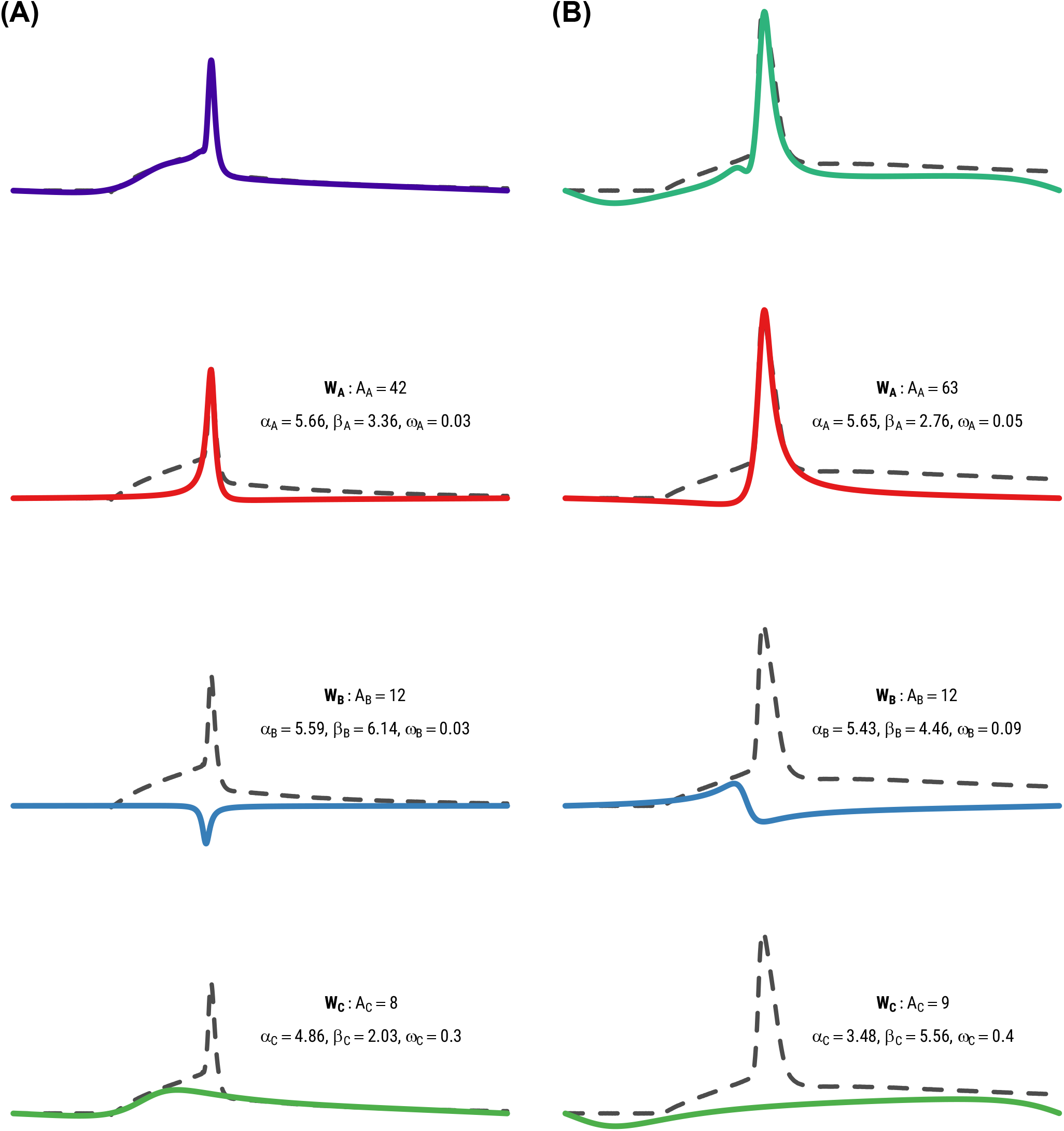
Fitted FMM model prediction and wave decomposition of a GABAergic AP (left) and a glutamatergic AP (right).

#### 2.1.2 Circular Principal Components Analysis (CPCA)

The CPCA is a procedure that generates a circular variable which gathers the maximum variability. A basic reference is Scholz (2007). Briefly, given a data base in matrix form, let e_1_ and e_2_ be the first two eigenvectors extracted with Principal Component Analysis (PCA, Hastie et al. (2009)). Consider the transformation in which the eigenvectors are projected onto the unit circle as follows:

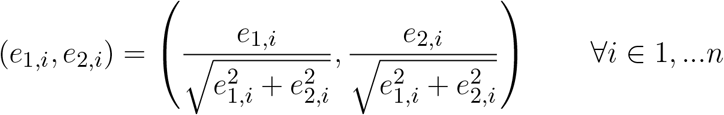

A circular order can be defined with *θ*_*i*_ = arctan 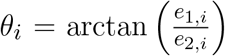, ∀*i* ∈ 1, …, *n*, which is called the first circular principal component.

#### 2.1.3 Machine Learning Supervised Methods

Various Machine Learning Supervised methods have been considered in the paper. The simple Linear Discriminant Analysis (LDA) method serves as benchmark for the results while, at the other extreme, the complex and “black box” methods Support Vector Machines of polynomial kernel (SVM) and Model Averaged Neuronal Network (AvNNet) methods have been considered. The former habitually achieves outstanding results in neuronal dynamics, as seen in (Teeter et al., 2018) among others. In between these two extremes, interpretable ensembles of decision tree methods have been used, particularly Random Forest (RF), which has been proved to attain great results without requiring precise hyperparameter tuning (as explained in Fernandez-Delgado et al. (2014)) and Gradient Boosting Decision Trees (GBDT), also capable of achieving outstanding results while not being as popular (Zhang et al., 2017). Some essential references to learn about these procedures are (Hastie et al., 2009) and (Izenman, 2008).

### 2.2 Dataset

The ACTD includes electrophysiological data of high temporal resolution of membrane potential from individual mouse recordings. A signal from each mouse neuron in the ACTD has been analysed; particularly the signal generated by the short square stimulus with the lowest stimulus amplitude that elicited a single AP. A small set of neurons that elicited two APs with the selected stimulus were initially discarded for the analysis. See (Allen Brain Institute, 2015) to learn about the stimulus types applied in the database. Each signal has been preprocessed and analysed according to the algorithm described shortly after.

A total of 1892 experiments, from mouse cells of 24 different Cre Lines, have been analysed. Beforehand, experiments from three Cre Lines were discarded as they did not have a sufficient sample size (less than 10 observations). The distribution of signals according to Cre Line is given in Table 1. Illustrated colours correspond to the different Cre Lines in all figures. To facilitate the reading, the appearance order of the Cre Lines in the table goes in accordance with the order proposed later in the paper. Throughout the manuscript, the characteristics of each Cre Line have been illustrated in two different ways: using median values and using representative neurons, selected from among the neurons in the Cre Line with the highest *R*^2^ that had all the extracted features between the 5th and 95th percentiles.

**Table 1.** Number of cells of each Cre Line by neuronal class.

| Cre Line | Total | Exc. | Inh. |
| --- | --- | --- | --- |
| Pvalb | 214 | 0 | 214 |
| Slc32a1 * | 27 | 0 | 27 |
| Nkx2.1 | 48 | 0 | 48 |
| Ndnf | 92 | 23 | 69 |
| Gad2 * | 19 | 0 | 19 |
| Htr3a | 159 | 10 | 149 |
| Sst | 120 | 2 | 118 |
| Nos1 | 67 | 6 | 61 |
| Oxtr * | 46 | 19 | 27 |
| Chrna2 | 70 | 25 | 45 |
| Chat | 67 | 0 | 67 |
| Vip | 122 | 19 | 103 |
| Ctgf | 59 | 55 | 4 |
| Tlx3 * | 40 | 40 | 0 |
| Sim1 * | 30 | 30 | 0 |
| Glt25d2 * | 10 | 10 | 0 |
| Ntsr1 | 67 | 62 | 5 |
| Esr2 * | 30 | 29 | 1 |
| Rbp4 | 87 | 86 | 1 |
| Scnn1a-Tg2 | 53 | 48 | 5 |
| Rorb | 173 | 150 | 23 |
| Scnn1a-Tg3 | 89 | 86 | 3 |
| Cux2 | 79 | 78 | 1 |
| Nr5a1 | 84 | 83 | 1 |

#### 2.2.1 Transcriptomic Features

In order to incorporate genetic information in the study, Cre Types in particular have been grouped into 8 transcriptomic genetic markers: Ndnf, Vip, Sst, Pvalb, L2-L4, L5, L6 and Non-neuronal. Some Cre Lines present in the current study were not present in the aforementioned paper. These Cre Lines without available transcriptomic features have been marked with an asterisk (*) throughout the paper.

### 2.3 Programming languages

The experimentation has been developed combining Python and R. Python has been used for data acquisition and transformation using the functions provided by Allen SDK (Allen Institute, 2015), while R fits the FMM models with the corresponding package available at the Comprehensive R Archive Network (https://cran.r-project.org/package=FMM) and analyses the results. The R packages (Venables and Ripley, 2002), (Karatzoglou et al., 2004), (Ripley, 2020), (Breiman et al., 2018), (Chen and Guestrin, 2016) and the auxiliary package for learning procedures caret (Kuhn, 2018) have been used to implement the Machine Learning procedures. Moreover, the libraries Shiny (Chang et al., 2020), Shinydashboard (Chang and Borges Ribeiro, 2018) and ggplot2 (Wickham, 2016) have been considered to develop a Shiny dashboard app.

### 2.4 Implemented algorithm

A flowchart of the preprocessing procedure and the estimation algorithm is depicted in Figure 2. First of all, in the preprocessing stage, the APs in the signal are extracted. Each AP segment is defined as [*t*_*S*_− 2*d, t*_*S*_ + 3*d*], with *t*_*S*_ denoting the time of the spike and *d* the time needed by the neuron to spike following the application of the stimulus. In real cases where the application time of the stimulus is unknown, a similar procedure can be applied preserving the uneven cut proposed. It is assumed that the segments to be analysed represent complete APs, in particular, *X*(*t*_1_) ≃ *X*(*t*_*n*_).

**Figure 2.**
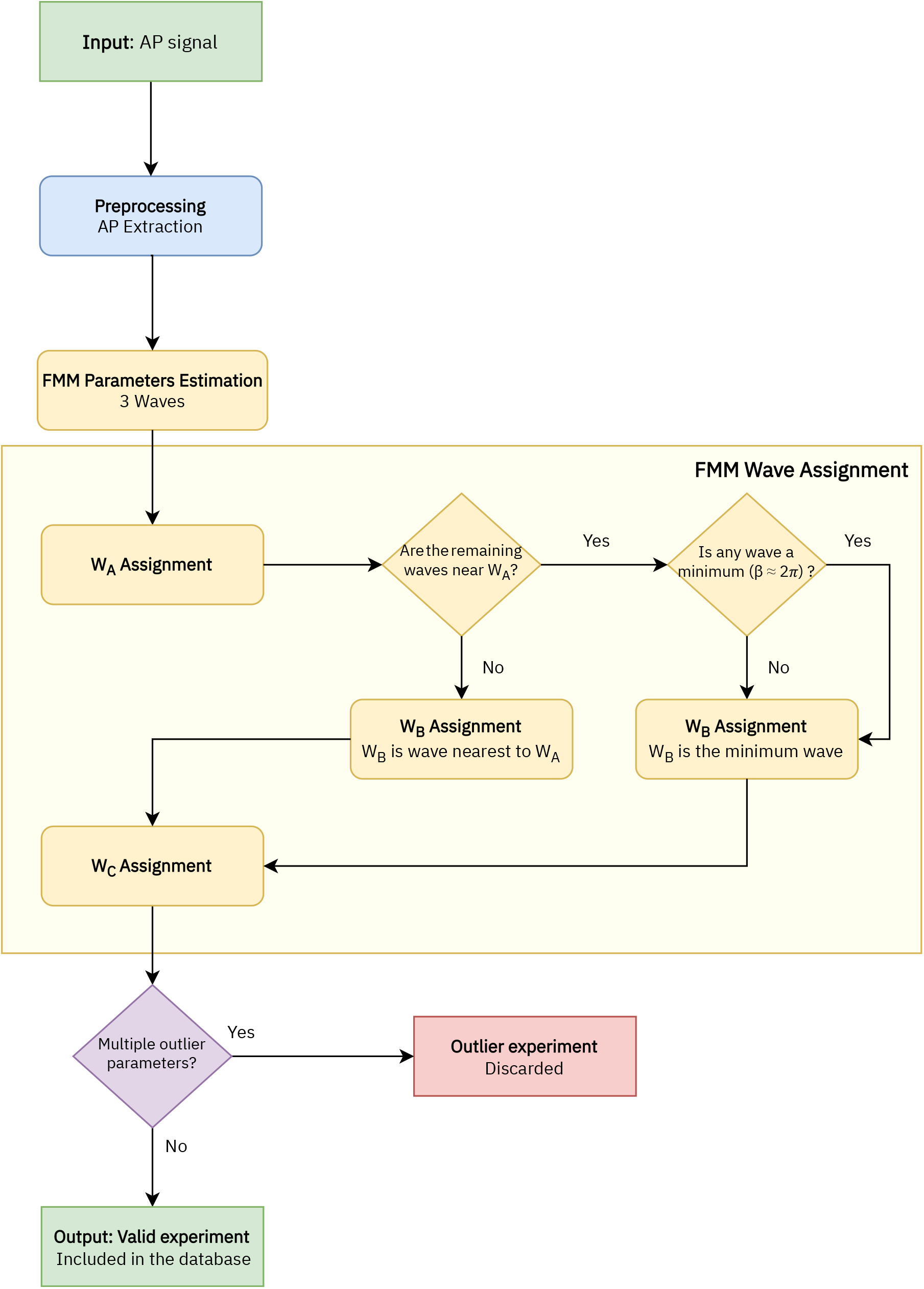
Flowchart of the implemented algorithm.

In a second stage, the parameters are estimated with the backfitting algorithm implemented in the package FMM of the programming language R, first presented in (Fernández et al., 2021). Three iterations of the backfitting algorithm are executed to extract the three waves. In each iteration, a single FMM wave is fitted to the residue of the previous iterations.

In the next stage the wave assignation is done. The proposed procedure is firstly presented in this paper and is specific to this study. Let the subscripts *i* = {1, 2, 3} denote the three estimated waves initially given by the backfitting algorithm. The labels *A, B* and *C* are assigned as follows:

1. *W*_*A*_ = *W*_*j*_*/j* = arg max_*i*=1,2,3_ *A*_*i*_.
2. Assuming the *W*_*A*_ = *W*_1_, *W*_*B*_ = *W*_*j*_*/j* = arg min_*i*=2,3_ *d*_*Ai*_, except in cases where |*d*_*A*2_− *d*_*A*3_| *<* 0.05 and min(*d*(*β*_2_, 2*π*), *d*(*β*_3_, 2*π*)) *<* 0.3, with *d*(*β*_*i*_, 2*π*) being the distance between the *β* parameter and 2*π*. In these cases *W*_*B*_ = *W*_*j*_*/j* = arg min_*i*=2,3_ *d*(*β*_*i*_, 2*π*).
3. Finally, *W*_*C*_ is the remaining wave.

The model is validated with the *R*^2^ statistic, which is the proportion of the variance explained by a model out of the total variance, as follows:

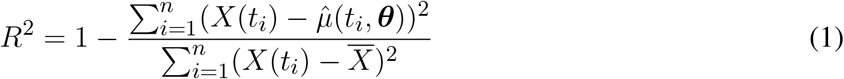

Finally, signals with multiple outlier values in significant parameters of the model related to the Cre Line distribution have been discarded.

## 3 RESULTS

### 3.1 FMM features for cell type characterization

The FMM model gives an accurate fit of the observed signals, the *R*^2^ global mean (standard deviation) being equal to 0.9868 (0.0066). GABAergic neurons are slightly better fitted as their mean *R*^2^ is 0.9903 (0.0054), while for glutamatergic neurons it is 0.9823 (0.0053). A Shiny app has been developed to illustrate the differences in the typical APs of the various GABAergic and glutamatergic Cre Lines. It can be accessed through https://alexarc26.shinyapps.io/medianapprofilebycreline/. The interface of the app, which is shown in Figure 3, consists basically of two parts: in the top half the median APs, along with their wave decomposition by Cre Line, are depicted, while the controls of the main figure are in the bottom half. These include the possibility of selecting the different Cre Lines of the database (up to nine different can be selected simultaneously), selecting just inhibitory or excitatory neurons, and selecting whether the wave sum or the parameters of the model should be plotted. A total of 40 cells have been discarded due to having multiple outlier values.

**Figure 3.**
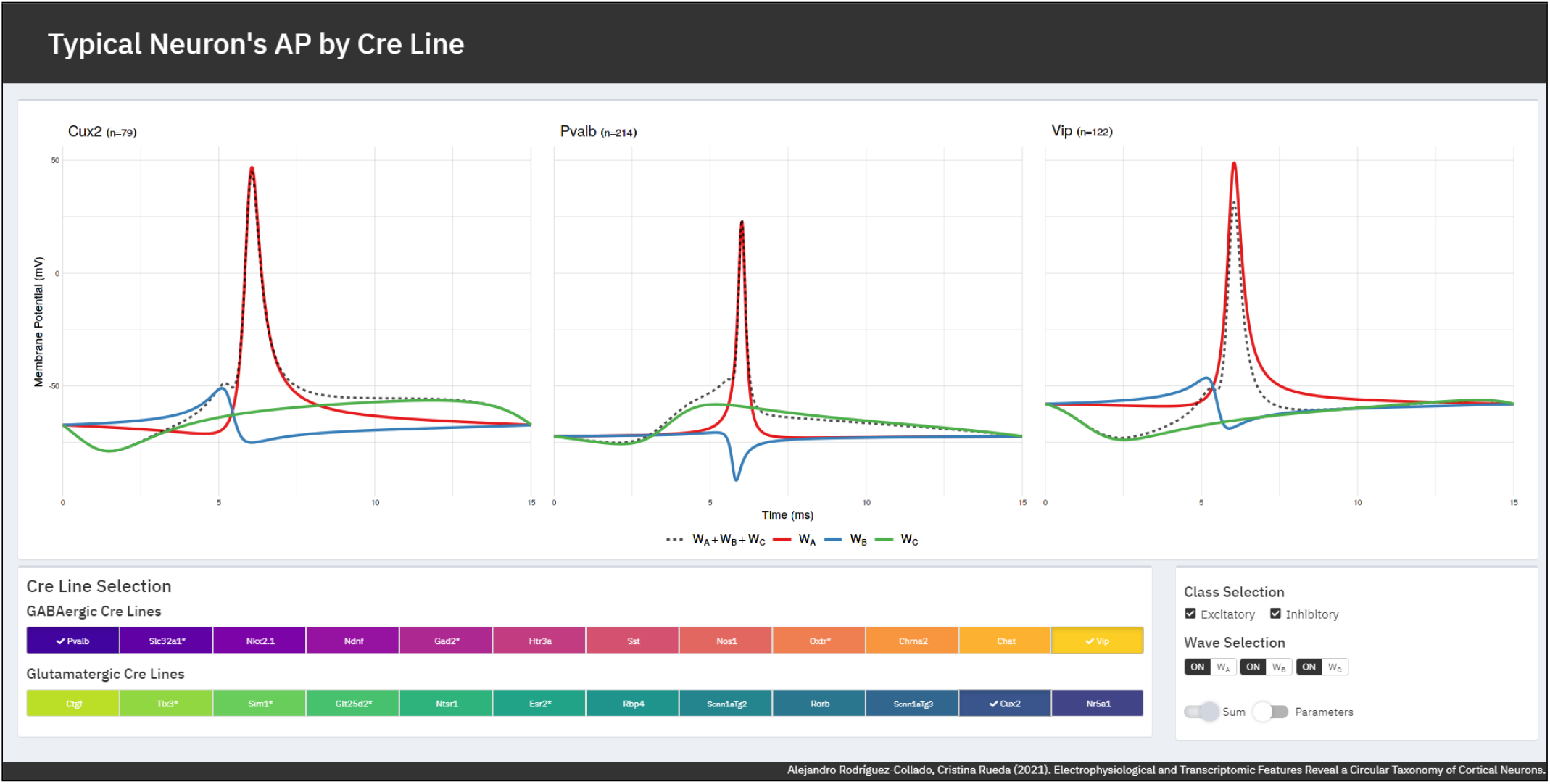
Typical AP by Cre Line Shiny app. In the top part the median APs along with its wave decomposition are depicted. The controls are in the bottom part of the interface.

The boxplots for the main parameters of the model by Cre Line are plotted in Figures S1-S5, in the Supplementary Materials. In these figures, the representative neurons of the Cre Line have been highlighted as stars. These plots illustrate the potential of various parameters to discriminate between the different Cre Lines such as *β*_*A*_, *ω*_*B*_ and sin(*β*_*C*_). The plots also show that the GABAergic neurons exhibit more variability in their electrophysiological features, as (Gouwens et al., 2019) points out.

Furthermore, the differences between Cre Lines are apparent not only in the time domain, but in the associated phase space. In Figure 4, the fitted FMM models and associated phase space of representative examples of GABAergic Cre Lines (A) and glutamatergic Cre Lines (B) are shown. Interesting differences can be seen both regarding the AP shape as well as the depicted trajectory in phase space.

**Figure 4.**
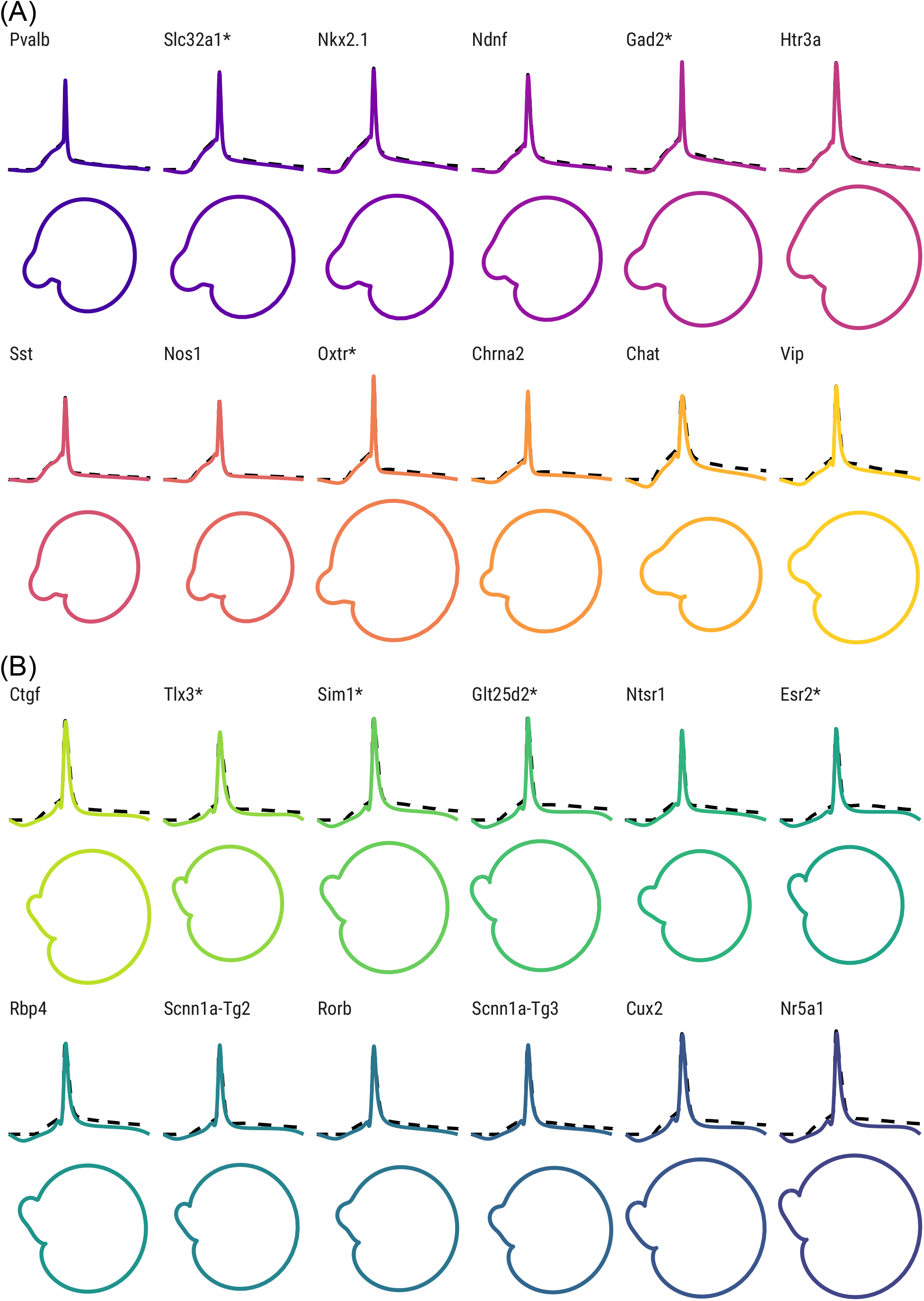
Fitted FMM model and associated phase space of representative examples of GABAergic Cre Lines (A) and glutamatergic Cre Lines (B).

### 3.2 Circular taxonomy

The taxonomy is defined at Cre Line level. For this purpose, the median values of the electrophysiological features by Cre Line have been used, along with the transcriptomic marker features. Due to notable distribution differences between the two feature sets, separate PCAs have been conducted. Firstly, the electrophysiological features PCA is conducted and two components are extracted (explained variance: 82.40 %). The correlation of the variables with the extracted components and the Cre Lines’ PCA projections and CPCA transformations are depicted in Figures S6 and S8 of the Supplementary Materials. The different Cre Lines are distinguished in the circular disposition and it is possible to tell apart most of the glutamatergic from GABAergic. This order is used later to place Cre Lines without having any transcriptomic features available in the taxonomy. The blank space between the Pvalb and Nr5a1 Cre Lines observed in Figure S8 corresponds to non-neuronal cells, unavailable in the studied data. Secondly, six components are extracted from the transcriptomic features as their explained variance was medium to low (93.47 %). The correlation of the variables with the extracted components and the Cre Lines’ PCA projections and CPCA transformations are depicted in Figures S7 and S9 of the Supplementary Materials. It is particularly relevant to note that the transcriptomic CPCA only distinguishes three groups of Cre Lines.

One final ensemble PCA is conducted with the extracted electrophysiological and transcriptomic components. The corresponding CPCA is shown in Figure 5, which is one of the main results of this study. The main novelty of the defined taxonomy is its circular topology, unlike the previous linear proposals. Moreover, for the first time to our knowledge, some Cre Lines have been located in a taxonomy. Nevertheless, a relevant difference with respect to other proposals, such as those of (Gouwens et al., 2019) and (Tasic et al., 2016), is that the Ndnf and Htr3a Cre Lines turn out to be similar to other GABAergic neurons, and not to non-neuronal cells. Further details of the final PCA results can be found in Figure S10 of the Supplementary Materials.

**Figure 5.**
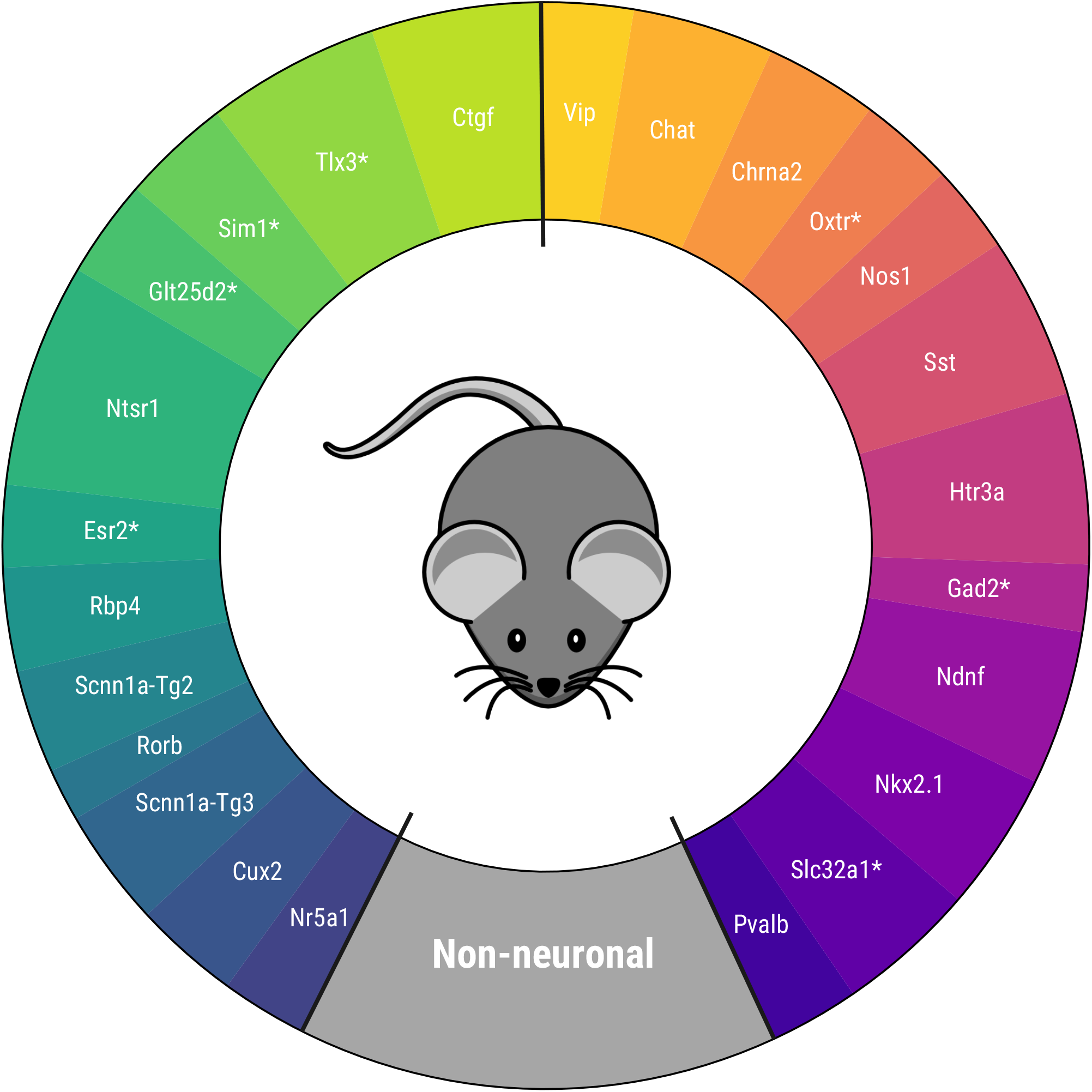
Proposed circular mouse cortical cell taxonomy.

The taxonomy is in agreement with others derived recently for mouse visual cortex neurons in several aspects. First, Cre Lines that have similar characteristics are kept together (Vip and Chat, Htr3a and Ndnf, Pvalb and Nkx2.1 among others) as in (Zeng and Sanes, 2017) and (Tasic et al., 2016). Second, the non-neuronal cell position between the GABAergic Pvalb Cre Line and the glutamatergic Nr5a1 and Cux2 Cre Lines -present in the upper layers of the visual cortex-coincides with the taxonomy of (Tasic et al., 2018). Within the glutamatergic neurons, the Ctgf and Ntsr1 Cre Lines-common in deeper layers- are the most similar in characteristics to the GABAergic neurons. In particular, this disposition is like those in (Gouwens et al., 2019) and (Tasic et al., 2016), after rearranging the results of the latter study, as can be seen in Figure S11 of the Supplementary Materials.

In order to validate the taxonomy, five transcriptomic-electrophysiological subclasses have been defined using Figure 5. These include four major GABAergic subclasses and one glutamatergic subclass specified in Table 2. In the next subsection, Machine Learning methods are used to discriminate these subclasses at neuronal level.

**Table 2.** Defined subclasses and Cre Lines composing them.

| 1. Pvalb+ |  |  | 2. Htr3a+ |  |  | 3. Sst+ |  |  |  | 4. Vip+ |  |
| --- | --- | --- | --- | --- | --- | --- | --- | --- | --- | --- | --- |
| Pvalb | Slc32a1* | Nkx2.1 | Ndnf | Gad2* | Htr3a | Sst | Nos1 | Oxtr* | Chrna2 | Chat | Vip |

| 5. Glutamatergic |  |  |  |  |  |  |  |  |  |  |  |
| --- | --- | --- | --- | --- | --- | --- | --- | --- | --- | --- | --- |
| Ctgf | Tlx3 * | Sim1 * | Glt25d2* | Ntsr1 | Esr2 * | Rbp4 | Scnn1a-Tg2 | Rorb | Scnn1a-Tg3 | Cux2 | Nr5a1 |

### 3.3 Cell-type classification

This classification problem, which is conducted at the cell level, has been addressed by other authors in many different ways, varying features and other factors such as the number, composition and definition of the subclasses or the selection of cells to be classified.

In this study, the FMM derived features have been considered. Specifically, 38 features have been used, including the basic parameters, peak and trough times -and their model values-, the explained variance of each wave and the distance between waves. Also, the cell’s reporter status, the origin layer and the applied stimulus amplitude have all been considered as predictors. Several Machine Learning methods were tested. Note that some classifiers assume that the predictors are euclidean, but *α* and *β* are circular parameters. The former and *β*_*A*_ can be considered euclidean as they take values concentrated in a small arc. However, sine and cosine transformations are applied to both *β*_*B*_ and *β*_*C*_.

All the classifiers except LDA had their hyperparameters tuned in a prior training-validation step. Afterwards, a ten-fold cross validation was performed on the tuned classifier to estimate its discrimination capacity. The dataset was divided into ten equally sized splits. In ten iterations, nine of the subsets were used to train the model, while the tenth serves as test. The discrimination capacity was evaluated in terms of the accuracy, percentage of observations correctly classified, and the kappa statistic, which measures the improvement over a random classification. A general overview of these matters can be found on (Hastie et al., 2009).

The classification problem was tackled in three different stages. The results can be seen in Table 3. In the first stage (A), the proposed classifiers were studied in the raw dataset, without discarding any observation or Cre Lines. More than 78 % of the cells could be correctly classified in their corresponding subclass by the AvNNet method; similar results were attained by SVM and RF. Observing the corresponding confusion matrix, it can be seen how the glutamatergic and Pvalb+ subclasses are most clearly discriminated, while more than half of the observations of the Htr3a+ subclass are misclassified.

**Table 3.** Cross-validated accuracy and kappa statistic of the different Machine Learning methods in the discrimination of the subclasses (left) and cross-validated confusion matrix of the best performer classifier -AvNNet in all cases-(right) in each of the defined stages.

|  | Accuracy | Kappa |
| --- | --- | --- |
| LDA | 68.8 % | 0.564 |
| RF | 77.1 % | 0.681 |
| GBDT | 76.5 % | 0.672 |
| SVM | 77.6 % | 0.687 |
| AvNNet | **78.3 %** | **0.697** |

|  |  | True class |  |  |  |  |
| --- | --- | --- | --- | --- | --- | --- |
|  |  | Pvalb+ | Htr3a+ | Sst+ | Vip+ | Glut. |
| Prediction | Pvalb+ | 88.2 % | 12.6 % | 11.6 % | 4.2 % | 0.9 % |
|  | Htr3a+ | 5.9 % | 48.2 % | 7.9 % | 11.1 % | 0.8 % |
|  | Sst+ | 4.5 % | 15.6 % | 59.4 % | 2.6 % | 1.0 % |
|  | Vip+ | 0.7 % | 9.3 % | 3.0 % | 64.0 % | 2.0 % |
|  | Glut. | 0.7 % | 14.4 % | 18.2 % | 18.0 % | 95.4 % |

**(B) Clean 5 Subclass Classification**
|  | Accuracy | Kappa |
| --- | --- | --- |
| LDA | 74.4 % | 0.645 |
| RF | 82.3 % | 0.755 |
| GBDT | 82.0 % | 0.751 |
| SVM | 83.2 % | 0.768 |
| AvNNet | **84.0 %** | **0.777** |

|  |  | True class |  |  |  |  |
| --- | --- | --- | --- | --- | --- | --- |
|  |  | Pvalb+ | Htr3a+ | Sst+ | Vip+ | Glut. |
| Prediction | Pvalb+ | 87.2 % | 12.2 % | 13.1 % | 4.7 % | 0.3 % |
|  | Htr3a+ | 6.9 % | 57.0 % | 8.8 % | 8.8 % | 0.3 % |
|  | Sst+ | 4.8 % | 17.3 % | 73.3 % | 4.1 % | 0.9 % |
|  | Vip+ | 0.7 % | 8.9 % | 2.0 % | 72.9 % | 1.3 % |
|  | Glut. | 0.3 % | 4.6 % | 2.8 % | 9.4 % | 97.2 % |

**(C) 4 Subclass Classification**
|  | Accuracy | Kappa |
| --- | --- | --- |
| LDA | 80.9 % | 0.706 |
| RF | 89.4 % | 0.841 |
| GBDT | 89.1 % | 0.837 |
| SVM | 91.2 % | 0.867 |
| AvNNet | **91.3 %** | **0.868** |

|  |  | True class |  |  |  |
| --- | --- | --- | --- | --- | --- |
|  |  | Pvalb+ | Sst+ | Vip+ | Glut. |
| Prediction | Pvalb+ | 93.1 % | 14.3 % | 5.3 % | 0.2 % |
|  | Sst+ | 6.5 % | 79.5 % | 4.1 % | 1.2 % |
|  | Vip+ | 0.0 % | 3.1 % | 82.4 % | 1.7 % |
|  | Glut. | 0.3 % | 3.1 % | 8.2 % | 96.9 % |

In the second stage (B), the GABAergic neurons that were not inhibitory and glutamatergic neurons that were not excitatory were discarded, leaving a total of 1704 observations. The accuracy is excellent for a five-class problem, with more than 84 % of the neurons being correctly discriminated by the AvNNet classifier. The second best result corresponds to SVM, followed closely by RF and GBDT, while LDA may be too simple for the task at hand. It is relevant to note that, in most of the cases, the misclassifications occurred between consecutive subclasses in the proposed circular taxonomy (i.e., Htr3a+ cells are mostly confused with Pvalb+ and Sst+, while Glutamatergic cells are misclassified with Vip+ cells). It seems that, at this stage, Sst+ and Vip+ subclasses are guessed correctly much better than in (A); the prediction of Htr3a+ has improved, but barely surpasses 55% of instances correctly classified.

Finally, in the third stage, the classification problem has been solved with 1304 observations, after discarding the observations from the Htr3a+ subclass as well as the observations from Cre Lines without transcriptomic features (marked with *). The results are outstanding, as more than 90 % of the observations are correctly discriminated in the proposed 4 classes. The best results are attained by the AvNNet and SVM classifiers. The Pvalb+ and Glutamatergic subclasses are well identified in more than 93 % of the cases, while Sst+ and Vip+ subclasses are approximately in 80 % of the cases.

The results of LDA clearly show evidence that the subclasses cannot be linearly discriminated. The RF and GBDT classifiers may have attained worse results than the “black box” methods in all the stages, but they offer interpretability in exchange, as feature relevance in the classification can be measured. In all the stages, the same features are highlighted as relevant. Particularly, *β*_*A*_ seems to be the most relevant feature, being at least 1.5 the relevance of the second most important feature in all cases. Other discriminant features are 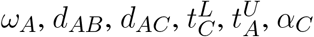, and the applied stimulus amplitude. The shape of the APs’ repolarization and depolarization phases captured by *W*_*A*_ seem to characterize the different subclasses.

## 4 DISCUSSION

In this paper, the FMM approach for electrophysiological feature extraction has been presented and used to describe a circular taxonomy in mouse cortical cells.

Relevant AP characteristics such as its width, amplitude, kurtosis, and skewness, among others, are represented by the FMM parameters. Furthermore, additional features standardly used in other studies can easily be defined in terms of the basic parameters, as has been done with the peak time and other features. Even more, the same set of parameters characterizes the phase space. As such it is not necessary to resort to additional feature sets, as is the case in (Gouwens et al., 2020). The FMM approach answers the necessities of *Feature Engineering* as the extracted parameters are interpretable and useful for classification and other data analysis problems.

A novel property to highlight of the proposed taxonomy is that it is circular. The latter addresses the need for neuronal types to be considered a continuum, discussed by many authors such as (Gouwens et al., 2020) and (Tasic et al., 2018). Previous proposals, being linear, consider cell types situated at the extremes to be opposite in terms of their characteristics. However, this does not reflect reality: cell types situated at the extremes habitually have a higher degree of similarity than the existing similarity between them and other types situated in the taxonomy’s centre. In fact, circular plots are used frequently to illustrate taxonomies. The taxonomy also follows the levels proposed by (Zeng and Sanes, 2017): at class levels, cells are either glutamatergic, GABAergic, or non-neuron while, at subclass level, GABAergic neurons can be either Pvalb positive, Vip positive, Sst positive, or Htr3a positive-Vip negative. Furthermore, at type level, the cells are classified according to the expressed Cre Line. In fact, some Cre Lines have been included in a taxonomy for the first time to our knowledge.

Despite some minor differences, the taxonomy’s Cre Line disposition proposal is in agreement with the literature about mouse visual cortex neuron types. Furthermore, the Cre Lines can be characterized using different FMM elements, such as its waves or parameters, that represent AP differences. In fact, the potential of the FMM parameters to discriminate glutamatergic neurons and the four major types of GABAergic neurons has been proved.

In short, the proposed taxonomy is hierarchical, continuous, easily reproducible, robust, and based on interpretable features.

A limitation of the presented study is that features have only been extracted from a single signal with a single AP generated from a short square stimulus. On the one hand, it remains to be seen if the application of the FMM approach on multiple signals of the same neuron could generate useful features. However, this question is up in the air as such authors as (Raghavan et al., 2019) suggest that the AP shape is independent of the applied stimulus. On the other hand, the use of the proposed method in multi-AP signals could be profitable, extracting other interesting features such as the interspike distance or the neuron’s firing rate.

As future work, the proposed taxonomy could be refined by using transcriptomic features at cell level, not only at Cre Line level. Also, the taxonomy should be validated further in other databases. The classifier methods presented in this work would profit from an increased sample size of each neuronal type and could probably discriminate better the different classes. In particular, the Htr3a+ subclass has turned out to particularly problematic to distinguish. However, other authors such as (Gouwens et al., 2018) have already remarked that electrophysiological features do not discriminate this neuron type particularly well.

## AUTHOR CONTRIBUTIONS

A.RC.: Developed the computational code for data acquisition and analysis, processed and analyzed data, wrote and revised the manuscript. C.R.: Conceived the aims, methodological proposal, conceptual design, wrote and revised the manuscript.

## DATA AVAILABILITY STATEMENT

The electrophysiological and transcriptomic data supporting the analysis of this study are available via the ACTD portal (http://celltypes.brain-map.org) and the Supporting Materials of (Tasic et al., 2016) respectively.

