## Supplementary Material for "Electrophysiological and Transcriptomic Features Reveal a Circular Taxonomy of Cortical Neurons"

#### 1 BOXPLOTS OF THE FMM PARAMETERS BY CRE LINE

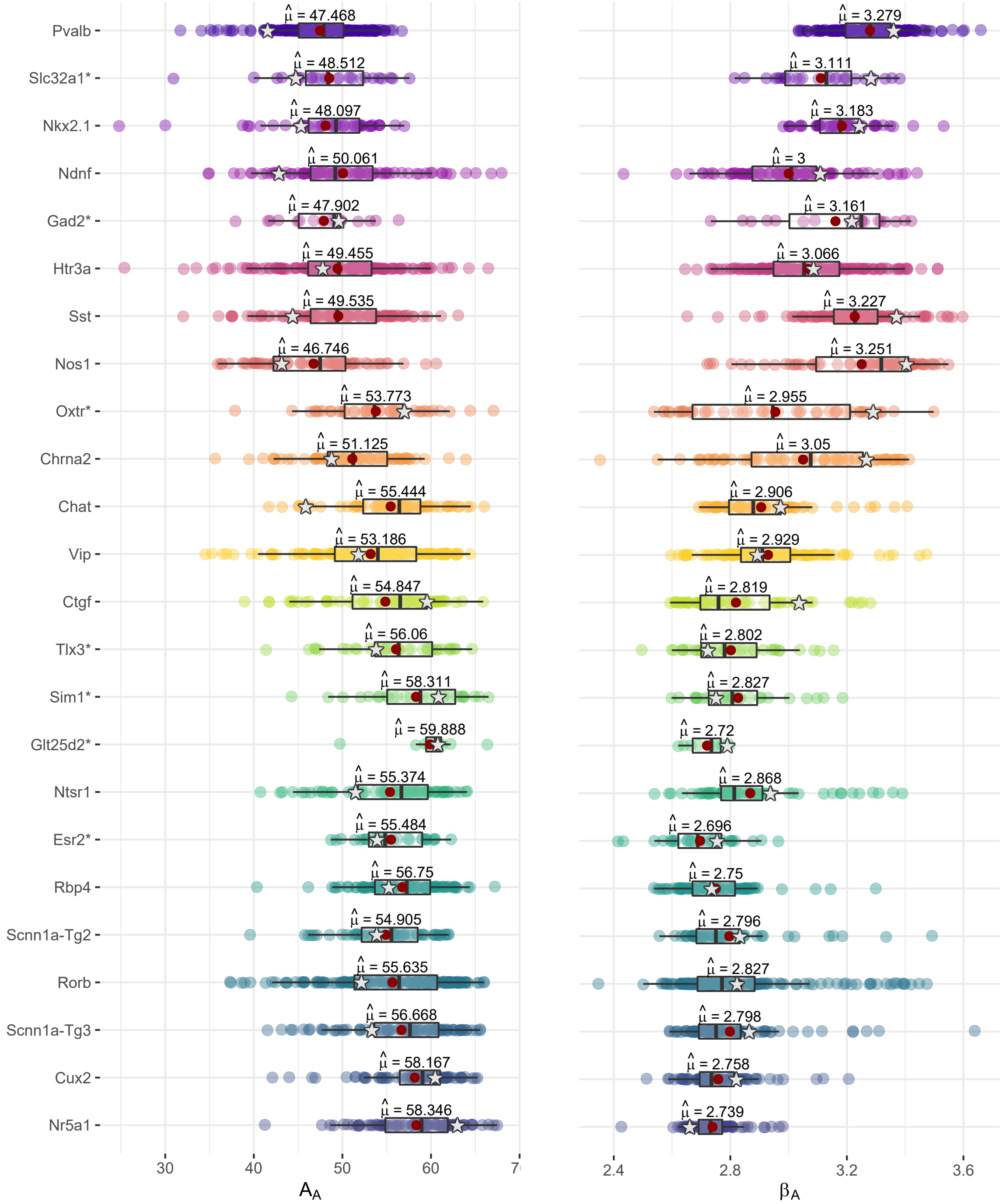


**Figure S1.** Distribution of the $A_A$ and $\beta_A$ parameters by Cre Line. Representative neurons used throughout the manuscript are highlighted as stars.

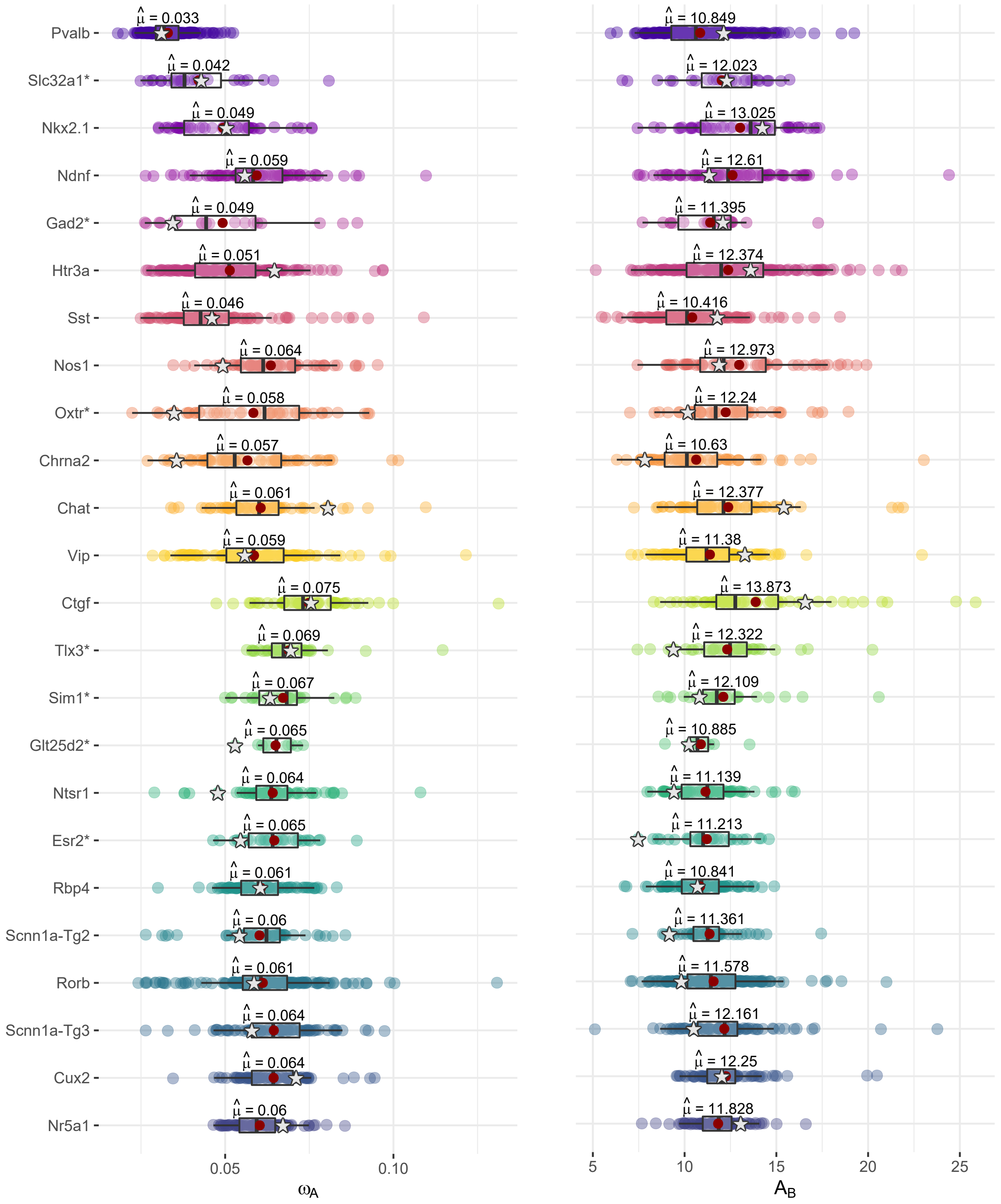


**Figure S2.** Distribution of the $\omega_A$ and $A_B$ parameters by Cre Line. Representative neurons used throughout the manuscript are highlighted as stars.

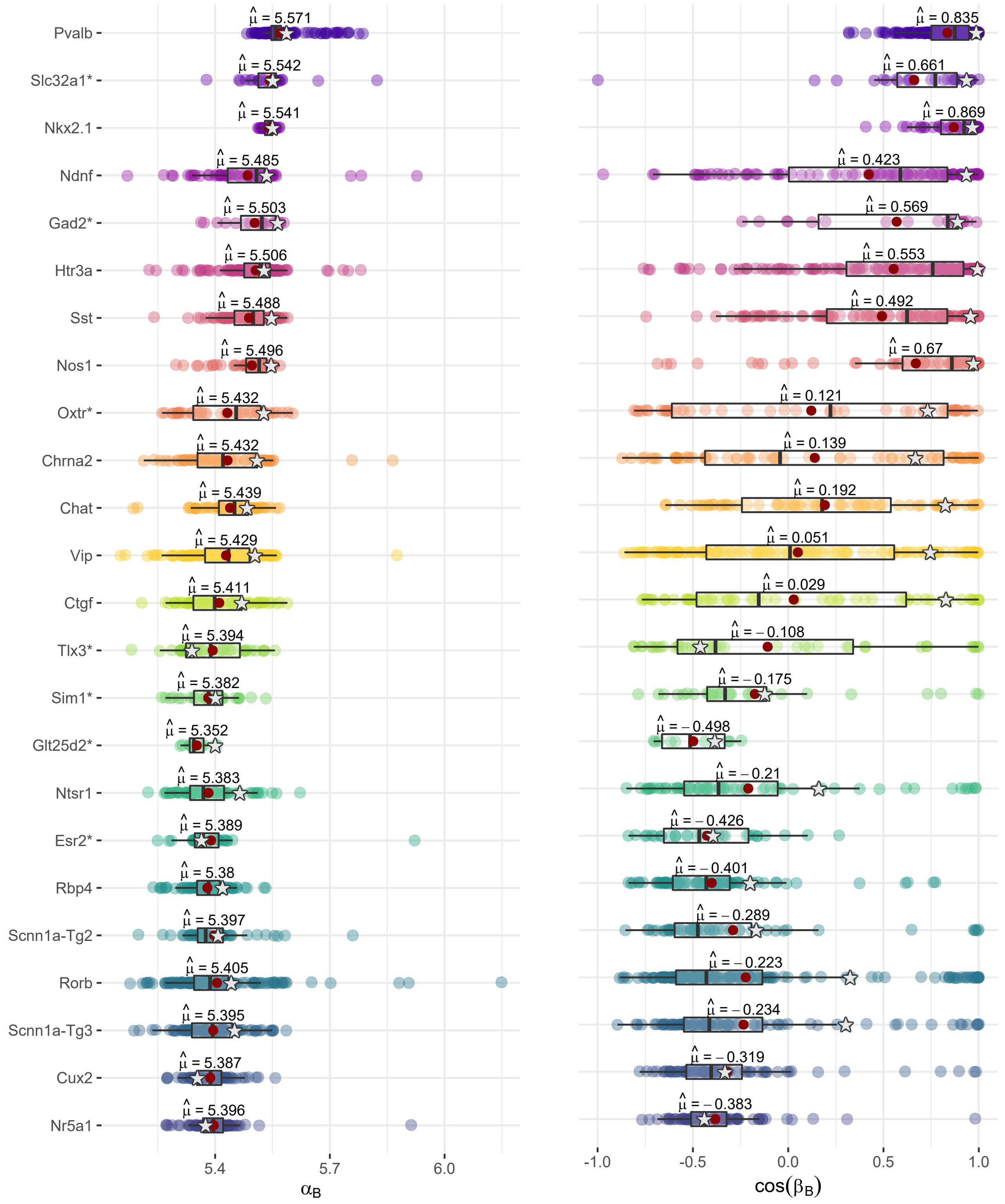


**Figure S3.** Distribution of the $\alpha_B$ and $\cos(\beta_B)$ parameters by Cre Line. Representative neurons used throughout the manuscript are highlighted as stars.

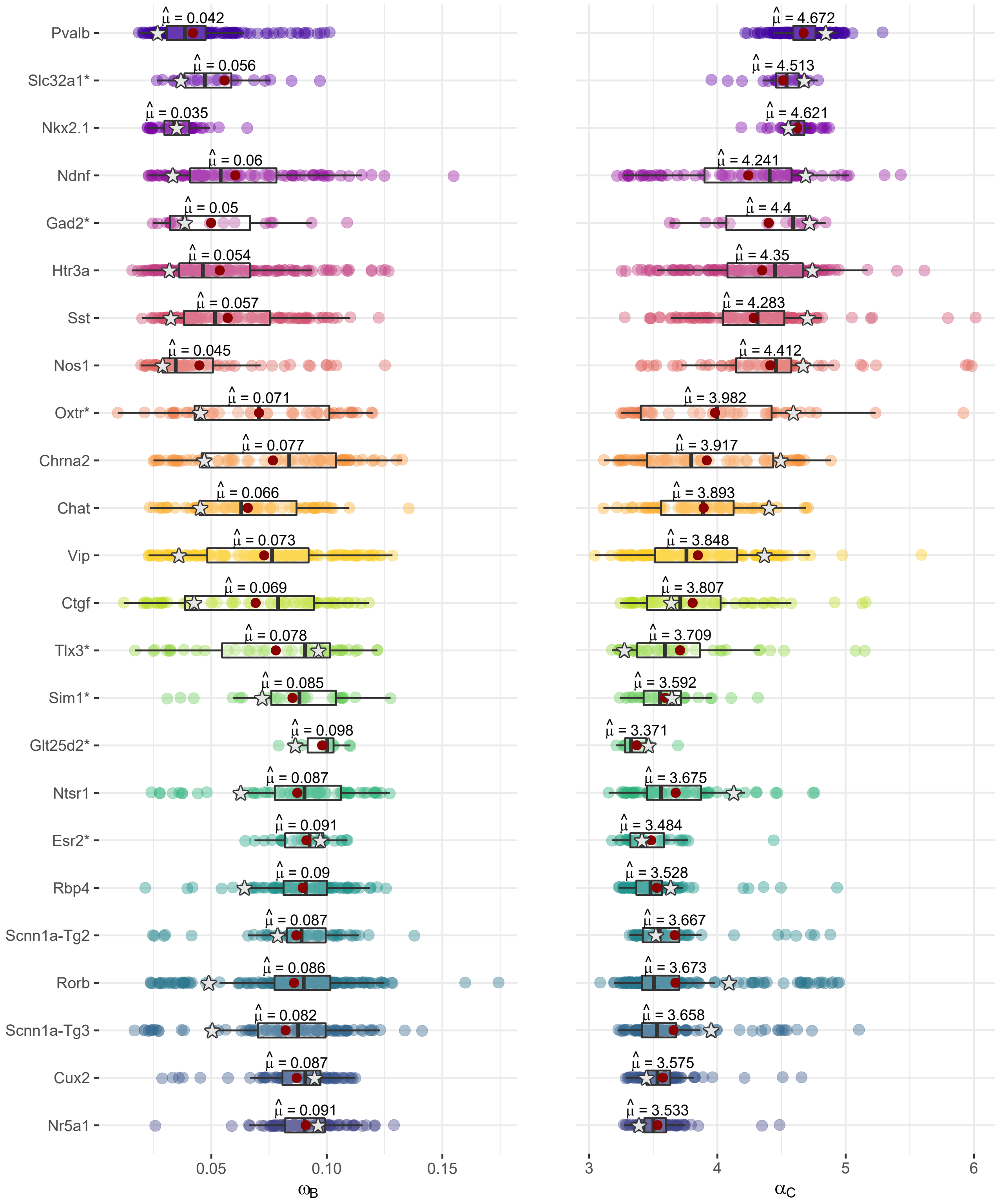


**Figure S4.** Distribution of the $\omega_B$ and $\alpha_C$ parameters by Cre Line. Representative neurons used throughout the manuscript are highlighted as stars.

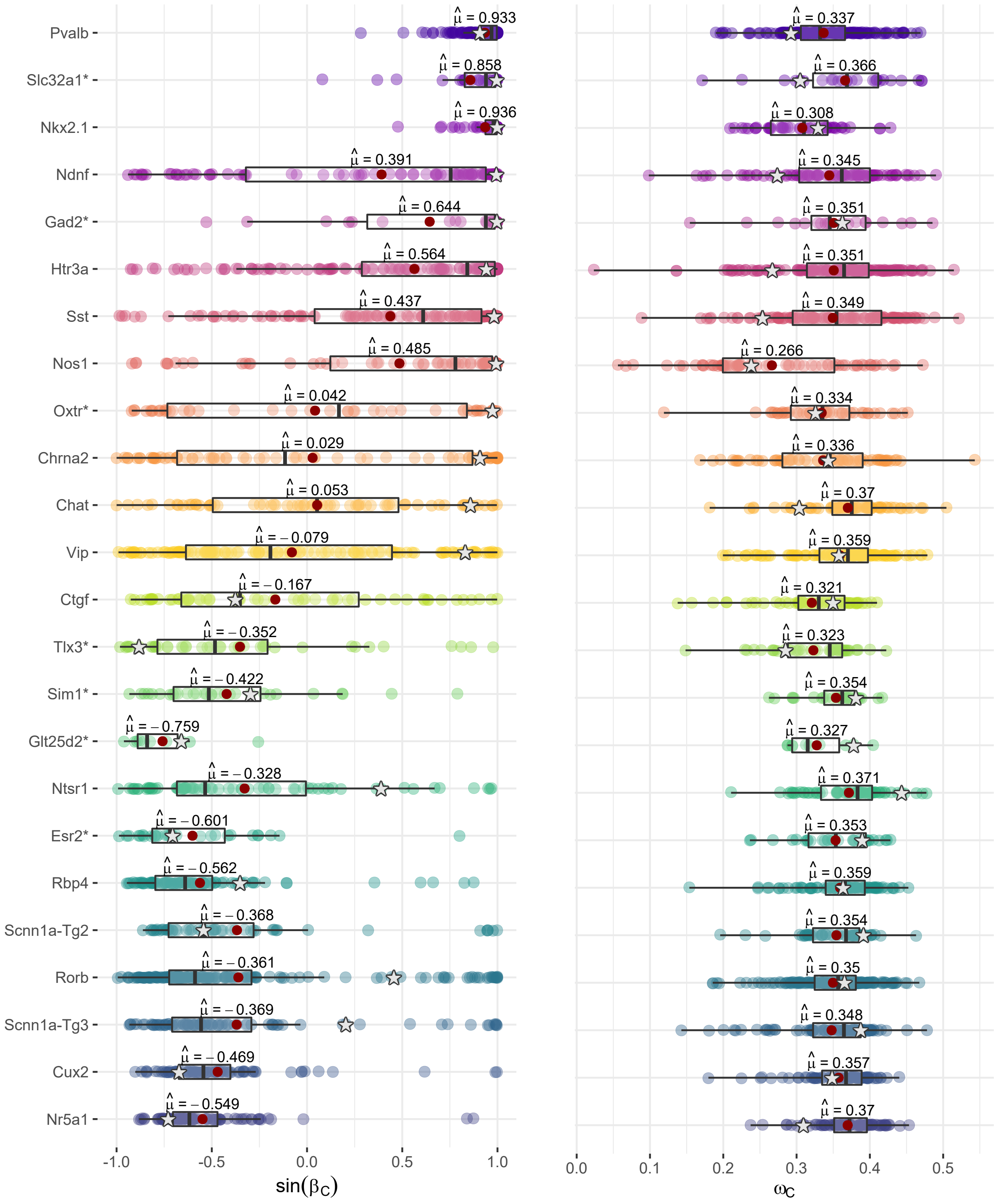


**Figure S5.** Distribution of the $\sin(\beta_C)$ and $\omega_C$ parameters by Cre Line. Representative neurons used throughout the manuscript are highlighted as stars.

#### 2 PRINCIPAL COMPONENT ANALYSIS (PCA)

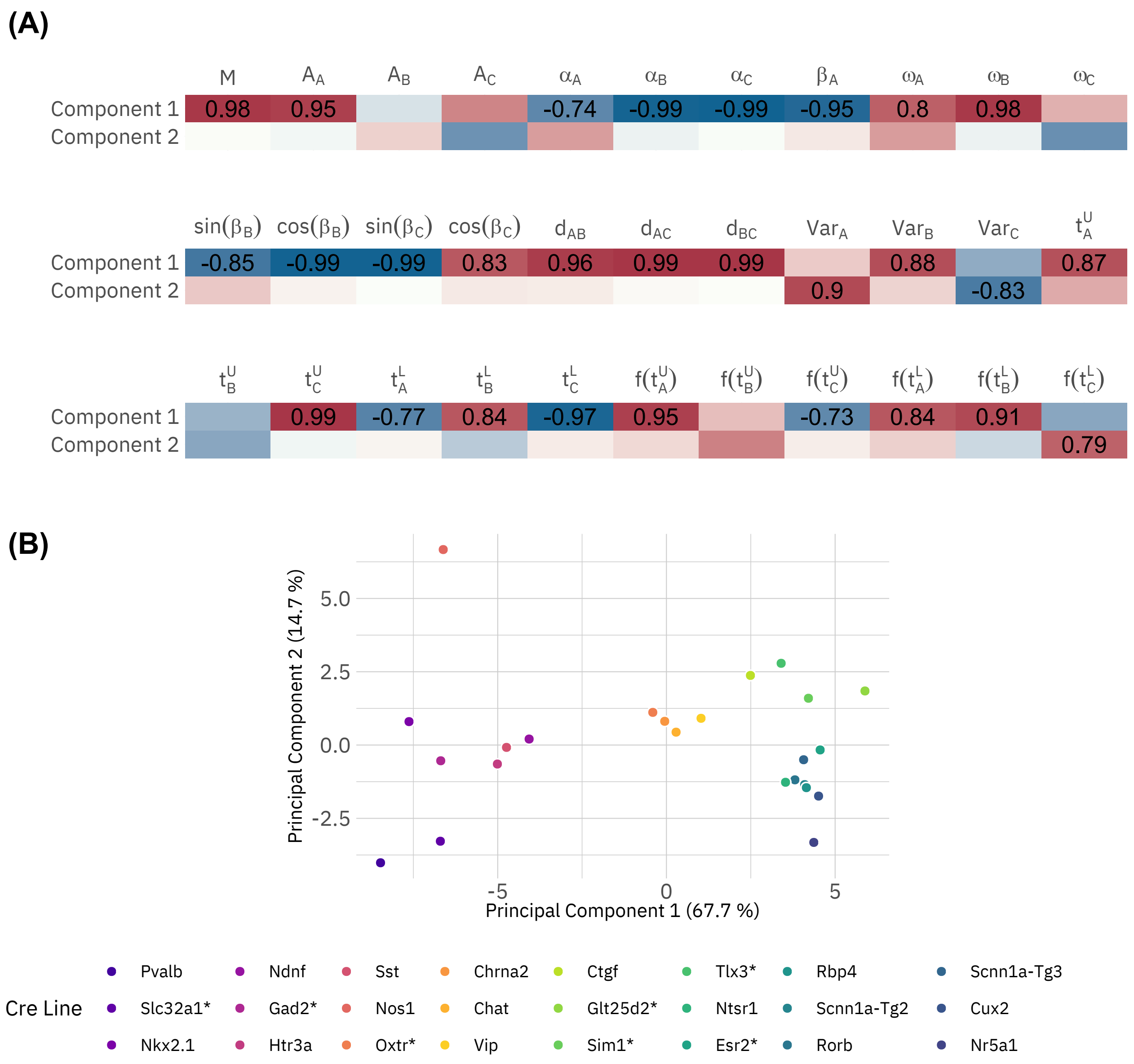


**Figure S6.** (A) Electrophysiological features correlation with extracted principal components. (B) Scores of the Cre Lines with the electrophysiological features PCA.

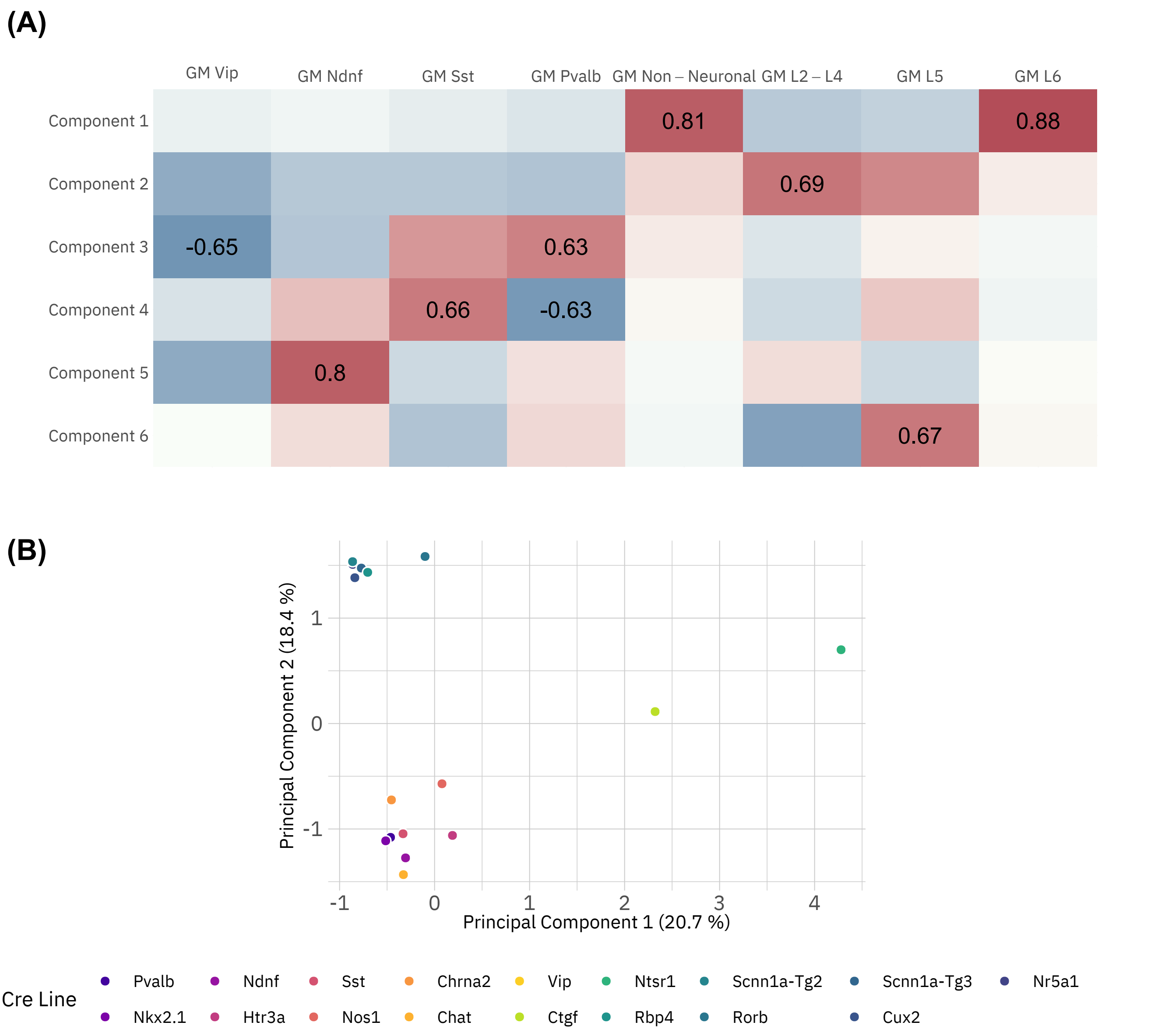


**Figure S7.** (A) Transcriptomic features correlation with extracted principal components. (B) Scores of the Cre Lines with the transcriptomic features PCA.

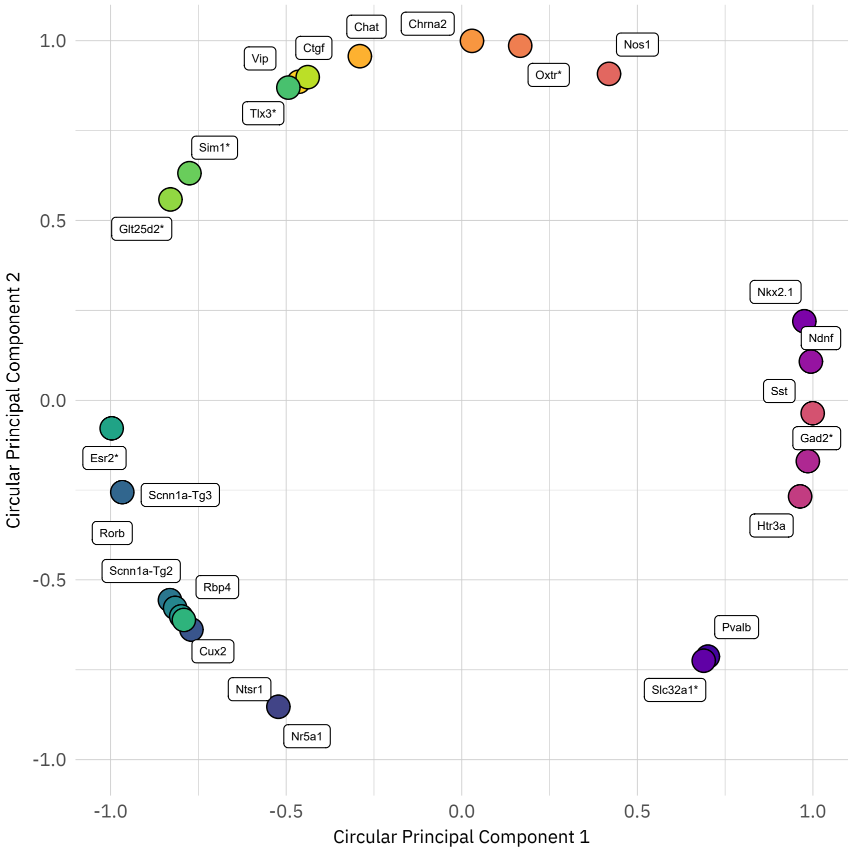


**Figure S8.** CPCA of the electrophysiological features.

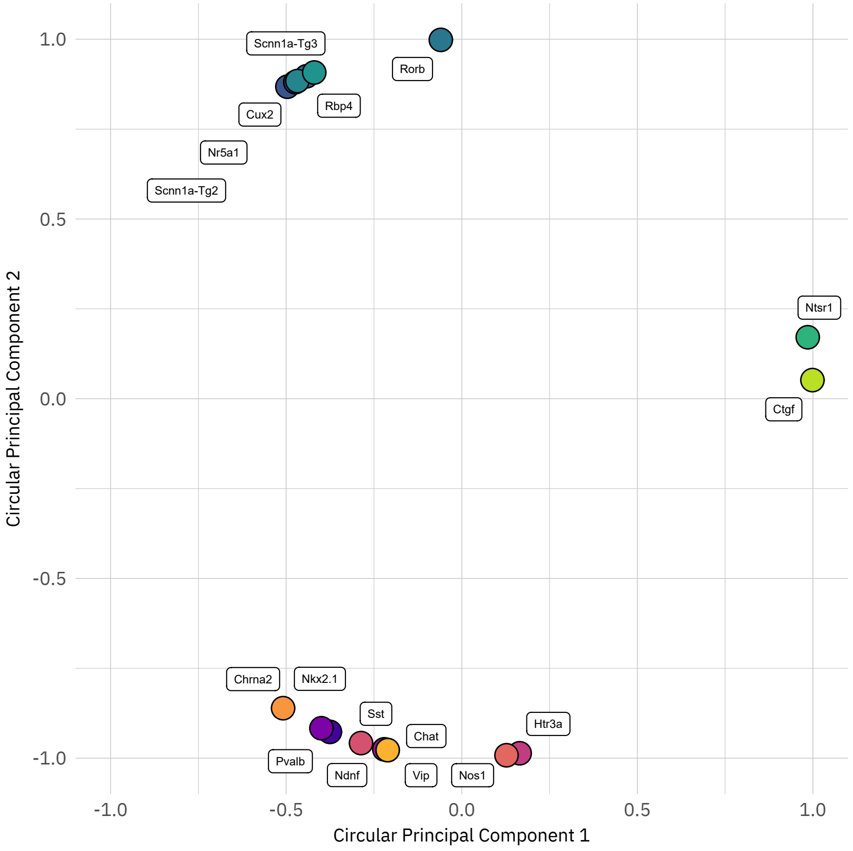


**Figure S9.** CPCA of the transcriptomic features.

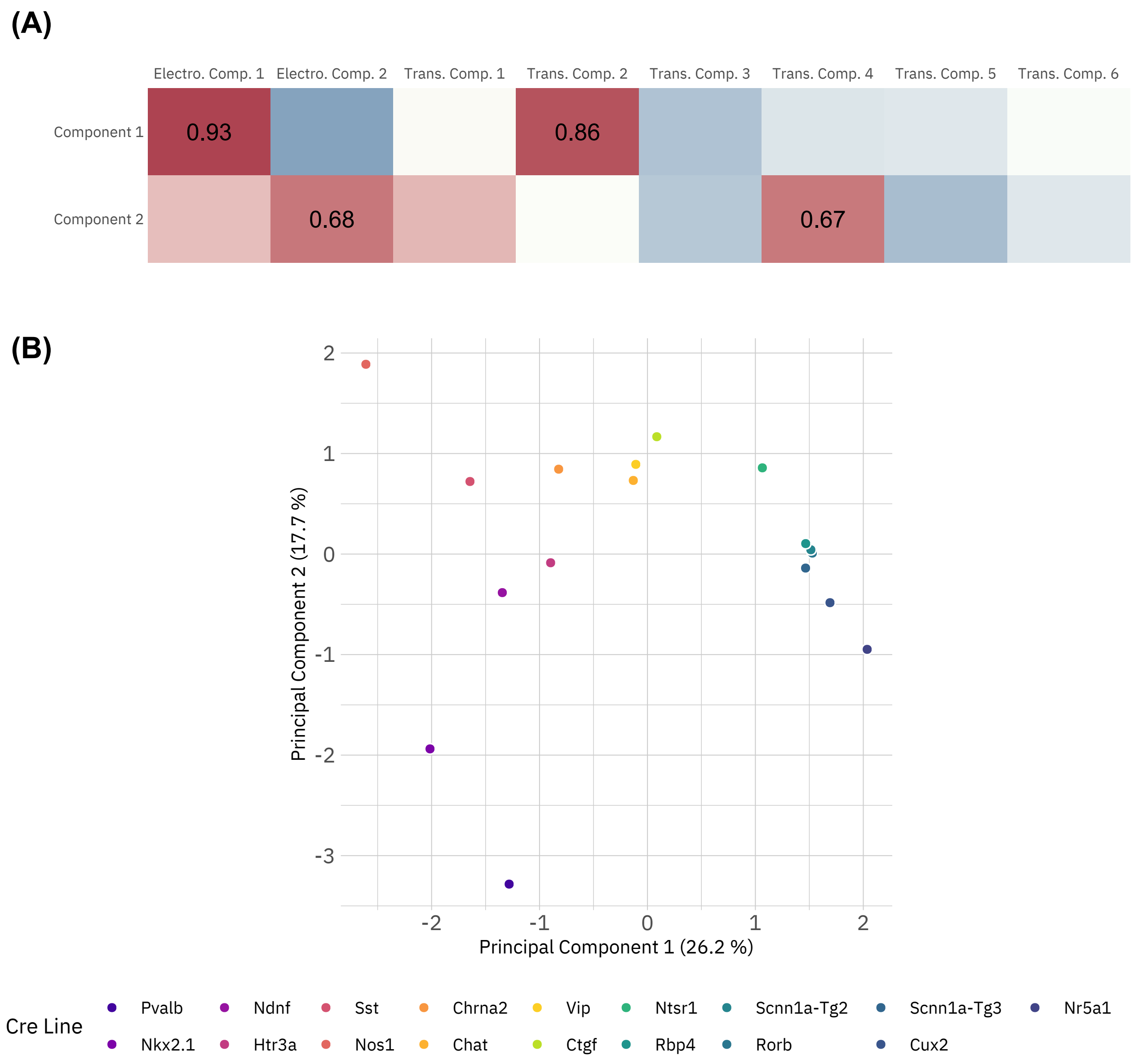


**Figure S10.** (A) Extracted electrophysiological and transcriptomic components correlation with extracted ensemble principal components. (B) Scores of the Cre Lines with the ensemble PCA.

### 3 TRANSCRIPTOMICAL NEURONAL MATRIX

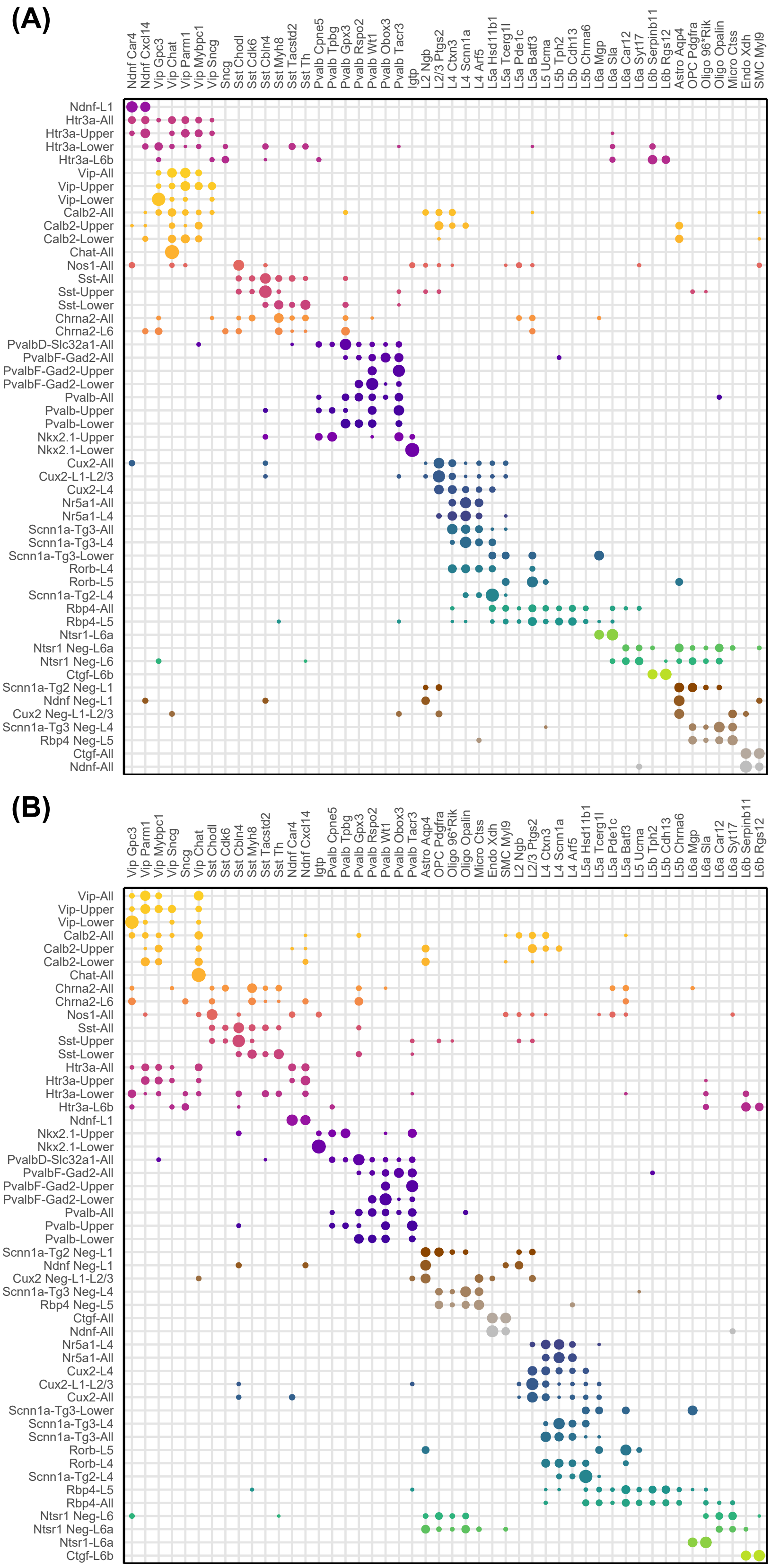


**Figure S11.** (A) Transcriptomic matrix from (Tasic et al., 2016). (B) Rearranged transcriptomic matrix according to the proposed circular taxonomy.
